# Social network cycle motifs and gut microbiome strain-sharing

**DOI:** 10.64898/2026.06.15.732275

**Authors:** Shivkumar Vishnempet Shridhar, George Iosifidis, Yanick Charette, Francesco Beghini, Nicholas A. Christakis

## Abstract

Microbes are commonly transmitted through human social interactions, yet little is known about how higher-order social network structures shape microbial circulation. Here, we analyze network cycles, i.e., closed loops of individuals, in both friendship networks and microbial networks, across 1,787 individuals from 18 isolated Honduran villages. Using strain-level resolution, we construct species-specific microbial networks, study their cyclic structure, and compare them with the social networks in the same population. Cycles were strongly over-represented relative to degree-preserving randomized networks in both social and microbial networks for most species, indicating that microbial transmission frequently occurs within recurrent and clustered groups of hosts. However, the overlap between microbial and social cycles varies substantially across species and individuals, and regression analyses identify a small subset of species whose cyclic sharing patterns are associated with social cycle participation. Notably, several anaerobic species show negative associations, suggesting reliance on repeated local exposures or shared environments only partially aligned with social ties.

Consistent with this, many species exhibit niche-like transmission pathways independent of social network structure. Together, these findings show that microbial sharing networks exhibit rich higher-order organization that only partially mirrors human social networks and that reveals that network cycles can provide a useful framework for understanding how repeated exposure might contribute to microbial spread.

## Introduction

Social interactions between hosts are a primary route for the transmission of both pathogens and commensal microbes.^1^ For the human gut microbiome, which is dominated by obligate anaerobes with limited survival capabilities outside their host, transmission typically requires close physical contact or shared living environments.^2,3^ Possible transmission can be identified via genetic analyses that identifies identical microbial strains between individuals, establishing what are known as “strain-sharing events”.^4^ While a significant portion of this exchange occurs within households,^2^ extensive strain-sharing also operates outside the households through direct friendships, and extends into indirect social relationships, such as second-degree (friend of friend) and third-degree (friend of friend of friend) connections.^1^ The ongoing circulation of gut microbes across these social structures may represent an important pathway for the persistence of beneficial commensal taxa in circumscribed human populations.^1,2,5–7^

However, traditional epidemiological frameworks often view transmission as a linear process or rely heavily on node properties, such as degree centrality (the total number of direct connections a person holds). While a high degree indeed exposes an individual to multiple independent neighbors, these exposures do not necessarily reinforce one another and frequently involve entirely distinct microbial strains. In contrast, sustained transmission and microbial persistence within a community may rely less on independent acquisitions from multiple disparate neighbors and more on repeated, short-range exposures within tightly connected groups. For some pathogens, such repeated exposures are required to establish a first infection or can increase disease severity^8–12^.

This clustered pattern of transmission is naturally captured by higher-order network topologies, and more specifically by network cycles. Defined as a closed path where the first and last nodes are identical, a *cycle* represents a loop of hosts among whom a microbial strain might continually circulate. The functional importance of network cycles is documented across diverse fields: in economic networks, cycles drive currency circulation and systemic financial health^13^, while in social networks, they can foster social capital and reciprocal exchange.^14,15^ Across these contexts, cycles capture feedback and reinforcement mechanisms that are entirely missed by analyzing only pairwise connections or open path motifs. In strain-level microbiome networks, cycles may provide a structural mechanism for repeated exposure over time, increasing the likelihood of strain reinforcement and long-term persistence within a population.

Here, we deploy a network-based framework to compute and analyze network cycles within microbial strain-sharing networks, assessing their relevance for host populations and individual bacterial microbes. Leveraging a comprehensive dataset of 1,787 individuals in 18 isolated Honduran villages, we construct strain-specific transmission networks for 639 microbial species. We first investigate which bacterial species commonly form cycles that may reflect circular transmission. Next, we evaluate the extent to which these microbial cycles structurally overlap with social networks reported by the humans. Finally, we implement a multi-layer network study, complemented with regression analysis, to compare cyclic profiles across species, providing insights into how higher-order human social structures may shape the baseline circulation of human microbiome.

## Results

### Construction of microbial and social networks

Using a cohort of 1,787 Honduran residents of 18 isolated villages, we study 639 species (see Methods for selection criteria) and their pattern of strain-sharing events. These species vary in presence and abundance, and have different strains. These strains can be shared between pairs of individuals and this can be quantified by assessing the similarity in genetic makeup between them, giving rise to strain-sharing ties^1^. These strain-sharing ties (or connections) between pairs of individuals result in microbial strain-sharing networks for each village and species (Figure 1A). Then, these strain-sharing ties can be analyzed in conjunction with social (friendship) networks, constructed independently using survey data (see Methods). Taken together, the 639 species yield a total of 4,240,736 intra-village ties between the 1,787 individuals (see Methods), out of a universe of 78 million ties that might conceivably exist among all individuals within each and every village. These create a total of 11,502 microbial networks across 639 species and 18 villages. All network ties were assumed to be undirected for analysis.

**Figure 1:**
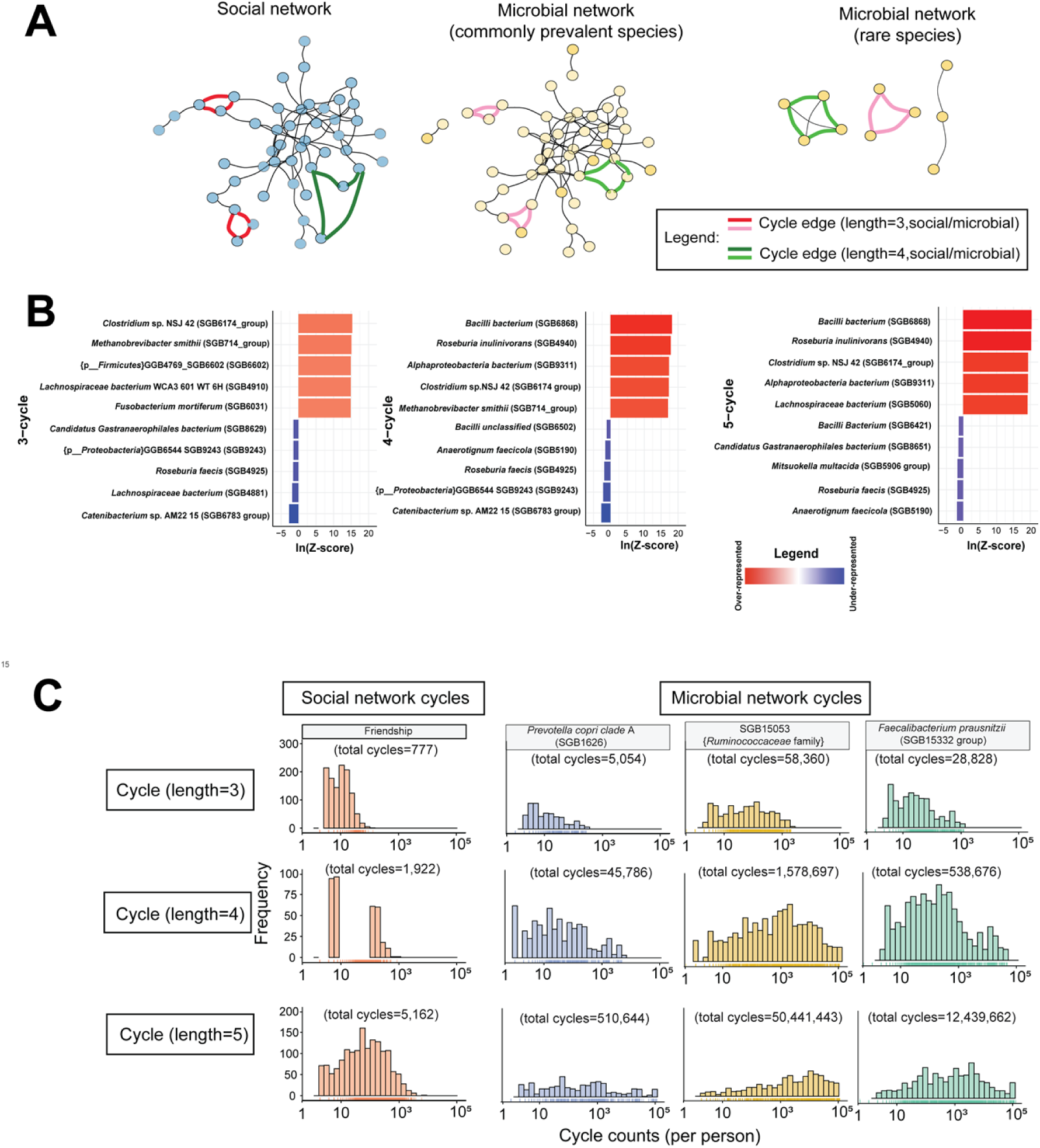
Cyclicity in microbial and social networks. (A) Social cycles, generated from social networks, and strain-sharing cycles, generated from microbial networks, are colored to show cycles, a closed-loop structure, in a demonstrative village, with each node representing a person residing in the village, and the edges showing either social or microbial interactions between individuals of the village. The microbial networks are shown for two species: a commonly abundant species - Escherichia coli (SGB10068) present in 1,765 individuals, and a rarely abundant unknown species from Clostridia family (SGB13986) present only in 492 individuals. The black lines indicate social connections (i.e., friendships) between pairs of individuals or the sharing of strains (i.e., microbial ties) between individuals. Green edges show cycles of length 4 and pink/red edges show cycles of length 3. (B) Enrichment of cycles in real network against permuted networks measured in Z-scores, with positive Z-scores representing over-representation of cycles, while negative scores show under-representation of cycles in real networks compared to null (permuted) networks. Top five and bottom five enriched species are shown for cycles of all lengths shown as ln(Z) where “ln” is natural logarithm. Some species appear across all studied cycle lengths. (C) Histograms showing individuals with varying participation in cycles in friendship networks (social network) and three most abundant species (microbial networks). Y-axis of the plot measures frequency of individuals in each bin, with the bins summing up to 1,787 individuals in all plots.

### Cyclicity of species

Social and microbial networks naturally contain cycles (closed loops) of varying lengths, defined by the number of individuals participating in each cycle. In the microbial strain-sharing networks, we identified 3,171,657 cycles of length 3; 61,563,292 cycles of length 4; and 1,621,373,460 cycles of length 5 across all species (Supplementary Table 1). We compared these networks with randomized counterparts preserving the same nodes, edges, and degree distributions. We found that 581, 475, and 516 out of the 639 analyzed species had significantly different null distributions for cycles of lengths 3, 4, and 5, respectively. Most species exhibited over-representation of cycles relative to the null model: 549 species (94.5%) for cycles of length 3; 422 species (88.8%) for cycles of length 4; and 479 species (92.8%) for cycles of length 5. In contrast, 32 (5.5%), 53 (11.2%), and 37 (7.2%) species had under-represented cycles of lengths 3, 4, and 5, respectively.

Among the most over-represented species were *Bacilli* bacterium (SGB6868), *Roseburia inulinivorans* (SGB4940) and Alphaproteobacteria bacterium (SGB9311) for cycles of length 4 and 5; and *Clostridium* bacterium *sp. NSJ 42* (SGB6174) and *Methanobrevibacter smithii* (SGB714) for cycles of lengths 3 and 4 (Figure 2B). Also, as expected, highly prevalent taxa (i.e., most common within this cohort), such as *Prevotella copri Clade A*, SGB15053 (*Ruminococcacae* family) and *Faecalibacterium prausnitzii* (SGB15332), showed consistent over-representation across all cycle lengths (permutation test p < 10^-25^; Fig. 1B, Fig. S1A).

**Figure 2:**
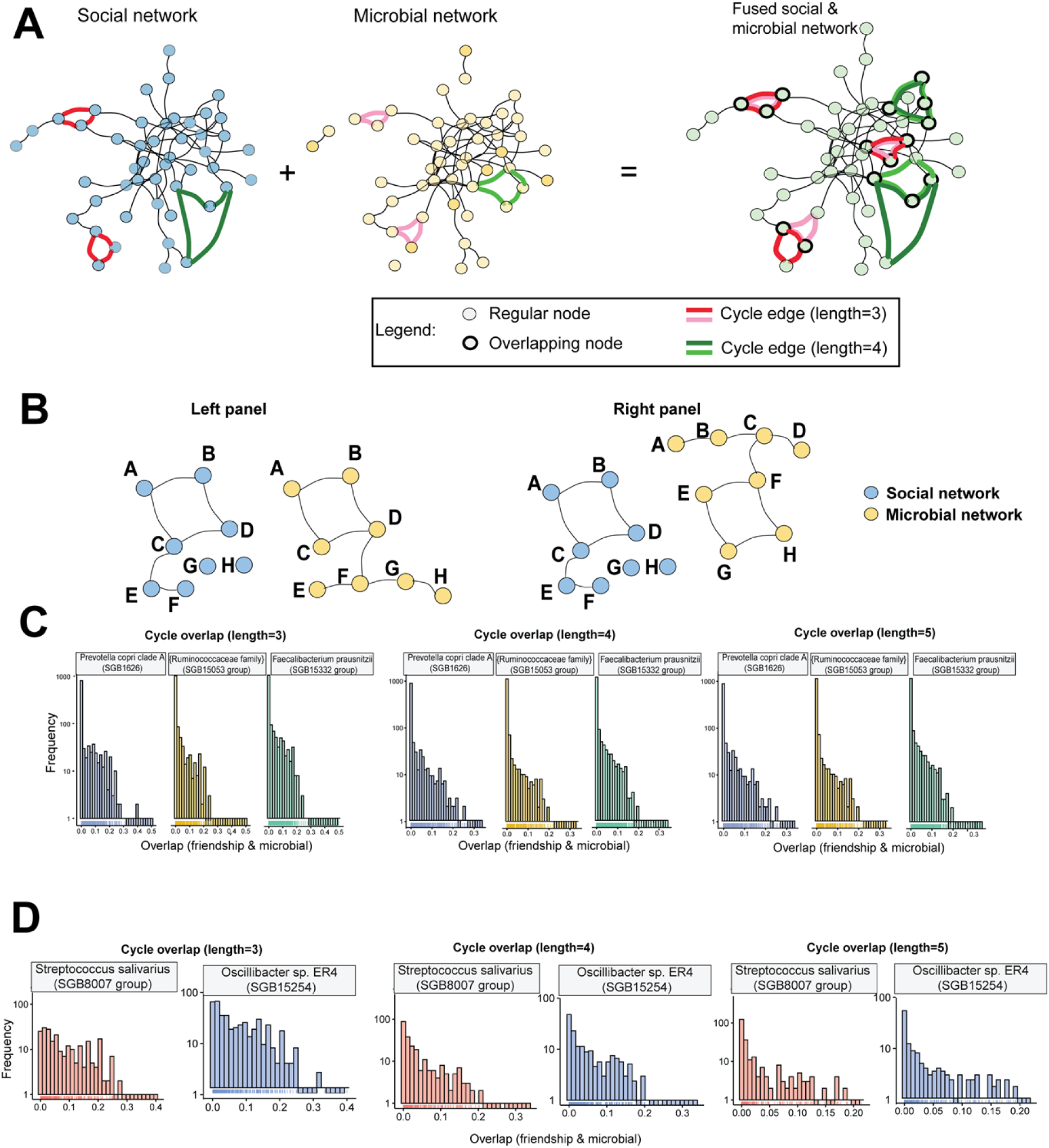
Overlap between social and microbial cycles. (A) Conceptual illustration of cycle overlap between social and microbial networks. Social cycles and microbial cycles are first identified independently and then fused to visualize overlapping nodes and edges. Overlapping nodes (with a thick black circle outline) represent individuals participating in both social and microbial cycles. Colored edges denote cycles of different lengths (red = length 3, green = length 4). The fused network shown corresponds to a representative village. (B) Another conceptual depiction, showing complete overlap in cycles in the left panel, against no overlap in cycles in the right panel across 8 individuals. (C) Distribution of the fraction of microbial cycles that overlap with social cycles, averaged per individual, for the three most abundant microbial species (*Prevotella copri* clade A, Ruminococcaceae SGB15053, and *Faecalibacterium prausnitzii*). Histograms illustrate the variability in social–microbial cycle overlap across individuals for cycle lengths 3, 4, and 5. (D) Same overlap analysis as in panel B, but shown for species identified as significant in the association model linking microbial cycle participation to social cycle participation (*Streptococcus salivarius* and *Oscillibacter* sp. ER4). These species were selected based on their statistical association with social network cycles rather than overall abundance.

Notably, several of the taxa with the strongest cycle enrichment, namely Bacilli bacterium (SGB6868), *Roseburia inulinivorans* (SGB4940) and *Faecalibacterium prausnitzii* (SGB15332), are anaerobic bacteria. Given their limited survival outside the host, the transmission of these bacteria is expected to depend on close contact and shared environments over short time scales. Their widespread occurrence therefore suggests repeated exposure within connected groups, making cycles a plausible structure of transmission. In addition, *Roseburia* and *Faecalibacterium* are fermentative anaerobes that produce butyrate and participate in cross-feeding interactions.^16,17^ These ecological dependencies may further favor repeated transmission within socially connected groups, consistent with the cyclic structures observed in their strain-sharing networks.

In contrast, *Roseburia faecis* (SGB4925) showed depletion across all cycle lengths, and *Catenibacterium sp. AM22-15* (SGB6783) was under-represented for cycles of lengths 3 and 4 (Figure 1B). To investigate this pattern, we examined the prevalence (i.e., number of carriers) for the species in conjunction with their strain-sharing connections. *Roseburia faecis* (SGB4925) was detected in 393 individuals and formed 983 edges, whereas *Catenibacterium sp. AM22-15* (SGB6783) was present in 1,619 individuals and formed 3,664 edges. By comparison, *Bacteroidales bacterium* (SGB2021) formed 3,996 edges despite fewer carriers (915 individuals). More broadly, cycle counts were strongly associated with prevalence, but substantial variation remained after accounting for it (Figure S2). For example, *Ruminococcaceae bacterium* (SGB14908) and the unclassified species SGB1664 (Prevotellaceae family) had similar prevalence (665 and 635 carriers, respectively), yet differed substantially in their number of cycles of length 3 (846 vs. 4,167), likely reflecting their different strain-sharing connectivity (2,250 vs. 7,529 edges). Indeed, cycle formation depends not only on species prevalence, but also on the density and topological arrangement of strain-sharing connections (p<0.05; Figure S2; Supplementary Table 1). Species with relatively fewer strain-sharing connections and cycles, such as *Roseburia faecis* and *Catenibacterium sp. AM22-15*, may therefore rely less on repeated transmission within social groups and more on environmental acquisition or broader dispersal processes.

To contextualize these patterns, we examined cyclicity in the human-reported social network. Using 4,324 undirected out-of-household friendship ties among 1,787 individuals, we identified 777 cycles of length 3; 1,922 of length 4; and 5,162 of length 5. As in microbial networks, cycles were significantly over-represented relative to permuted networks (p < 10^-25^; Figure S1C), consistent with prior evidence that social networks exhibit cycle structures.^15^ Z-scores from permutation tests were large (Methods), and are reported as ln(Z) for comparability: 5.15 (length 3), 5.22 (length 4), and 15.22 (length 5).

Overall, both social and microbial networks exhibit strong cyclic structure relative to their randomized counterparts, suggesting that repeated interactions within socially connected groups contribute to microbial transmission. Because cycle counts scale with population size, subsequent analyses focus on per-individual cycle participation (cyclic centrality).

### Heterogeneity in cycle participation across individuals and species

We next shift our focus from network-level properties to individual-level participation in social cycles and strain-sharing cycles. To quantify heterogeneity in cycle participation, we use k-cycle centrality^15,18^, defined here as the number of cycles of length *k* involving a given individual, normalized by the total number of cycles in the village. Using this metric, we examined the extent to which participation is concentrated in a small subset of individuals, by fitting discrete power-law distributions. Model parameters and the lower bound *x_min_* were estimated using standard procedures, and goodness-of-fit was assessed using a bootstrap-based Kolmogorov–Smirnov (KS) test with 500 simulations (see Methods). Bootstrap p-values greater than 0.1 indicate the power-law model is a statistically plausible description of the upper tail of the distribution.

Participation in social network cycles exhibited a heavy-tailed distribution consistent with a power-law tail for cycles of length 3 and 4 (power-law *p* = 0.32 for length 3, *p* = 0.48 for length 4). For cycles of length 5, the tail was better approximated by a log-normal distribution (*p* = 0.99), indicating that, while most individuals participate in relatively few social cycles, a small number participate in very large numbers of cycles. Overall, cycle participation in social networks was strongly right-skewed, spanning several orders of magnitude from only a few cycles to thousands per individual.

We next examined the microbial strain-sharing networks cycle distribution. Among the most prevalent species, several microbial networks also displayed heavy-tailed distributions of cycle participation. For example, *Streptococcus salivarius* showed a power-law–like tail for cycles of length 3, while *Faecalibacterium prausnitzii* and *Oscillibacter sp. ER4* (SGB15254) exhibited heavy-tailed participation patterns across multiple cycle lengths. In contrast, other prevalent species, such as *Prevotella copri* Clade A did not conform to either power-law or log-normal tail models. Other species, such as SGB15053, showed distributions more consistent with log-normal behavior (Figure 1C).

Together, these results indicate that cycle participation in both social and microbial networks is highly concentrated among a relatively small subset of individuals. Although microbial species differed in the precise form of their tail behavior (Supplementary Table 3), many exhibited strongly right-skewed participation patterns like those observed in social networks. These similarities suggest that cyclic organization in microbial strain-sharing networks may partially reflect the structure of underlying social interactions. And, further, the heterogeneity across species indicates that this relationship is unlikely to be uniform across microbes and may depend on species-specific transmission mechanisms, colonization dynamics, variations in environmental persistence, and metabolic interactions within the microbial communities.

### Relationship between social and microbial cycle centrality

Motivated by the shared heavy-tailed structure of social and microbial cycle participation, we next examined whether individuals occupying highly cyclic positions in social networks also participated more strongly in cyclic structures within microbial strain-sharing networks. Previous work has shown that microbial strain-sharing predicts social relationships between pairs of individuals.^1^ Here, we extend this framework beyond pairwise relationships to cycles, and also test whether this relationship between social and microbial cycle participation varies across species.

To that end, we measured cycle centrality for each individual in both the friendship network and species-specific microbial networks (see Methods). We then modeled the association between social and microbial cycle participation across species and cycle lengths. Models included village as a random effect to account for shared environmental and social structure within villages, while controlling for node degree as a proxy for local connectivity. Following our previous association framework^7^, we used beta regression with microbial cycle participation as the outcome variable. To further control for cyclic structures arising purely from network density, we performed permutation tests using randomized networks generated separately for each species-village combination while preserving node degree distributions (see Methods).

Overall, significant associations between social and microbial cycle participation were observed for only a subset of species, indicating the relationship between social and microbial cyclicity is species-specific rather than universal. For cycles of length 3, significant associations were identified for *Streptococcus salivarius* (SGB8007 group), SGB1883 (*Bacteroidaceae* family), and Candidatus *Gastranaerophilales* bacterium (SGB8609). For cycles of length 4, significant associations were detected for *Streptococcus salivarius*, SGB15053 (Ruminococcaceae family), *Oscillibacter* sp. *ER4* (SGB15254), SGB1667 (*Prevotellaceae* family), and SGB1883. For cycles of length 5, significant associations were observed for *Streptococcus salivarius*, SGB15053, *Oscillibacter* sp. *ER4*, SGB1667, SGB1883, and SGB15085 (*Ruminococcaceae* family).

Although both positive and negative associations were observed, only a limited subset remained significant after correction for multiple testing.

Notably, microbial cycle centrality in the *Streptococcus salivarius* (SGB8007 group) network was negatively associated with social cycle centrality across all cycle lengths examined (β₃ = - 0.28, β₄ = −0.09, β₅ = −0.03; Supplementary Table 4). This is primarily an oral microbe, although it can also colonize the digestive tract, and has been linked to dietary and environmental acquisition pathways.^19,20^ Its weaker alignment with social cycle structure may therefore reflect transmission routes that depend less strongly on repeated close social contact and more on environmental exposure, including aerosolized droplets.^21^

Similarly, for cycles of lengths 4 and 5, social and microbial cycle participation were negatively associated for SGB15053 (*Ruminococcaceae* family; β₄ = −0.025, β₅ = −0.007) and *Oscillibacter* sp. ER4 (SGB15254; β₄ = −0.03, β₅ = −0.01). Members of the *Ruminococcaceae* family are known for fiber degradation,^22^ whereas *Oscillibacter* species have been associated with anti-inflammatory functions.^23^ Both taxa are anaerobic, and their weaker correspondence with social cycle structure may indicate that host physiological or dietary factors contribute more strongly to their colonization and persistence than repeated transmission within social network cycles.

### Overlap between social and microbial cycles

So far, we have analyzed cycle structure separately in social and microbial strain-sharing networks. However, both networks are defined over the same set of 1,787 individuals, allowing us to examine the extent to which cyclic structures in the two settings involve actual overlapping groups of individuals (i.e., not just similar metrics of k-cycle centrality). We quantified the overlap between social and microbial cycles by measuring the similarity between the sets of individuals participating in cycles across the two network types (Figure 2A–B). In particular, we used the Jaccard index to compare the sets of alters participating in cycles with a given ego in the social network and in each species-specific microbial network. Overlap values can thus range from 0 (no shared participants between social and microbial cycles) to 1 (complete overlap of cycle participants across modalities) (see Methods). This analysis was performed separately for each species and paired with social network for cross-modal analysis.

We first examined how this overlap was distributed across individuals, and whether these distributions differed across microbial species. For cycles of length 3, *Oscillibacter* sp. ER4 (SGB15254) and *Faecalibacterium prausnitzii* (SGB15316 group) exhibited strongly right-skewed overlap distributions consistent with log-normal behavior, indicating that overlap between social and microbial cycles was concentrated among a relatively small subset of individuals. In contrast, *Streptococcus salivarius* (SGB8007 group), *Prevotella copri* Clade A (SGB1626), and SGB15053 (Ruminococcaceae family) showed substantially less skewed overlap distributions, suggesting more evenly distributed overlap across individuals (Figure 2B–C; Supplementary Table 5).

For cycles of length 4, overlap distributions remained heterogeneous across species. *Streptococcus salivarius*, *Prevotella copri* Clade A, and *Oscillibacter* sp. ER4 displayed heavy-tailed overlap distributions consistent with log-normal behavior, whereas *Faecalibacterium prausnitzii* and SGB15053 exhibited overlap distributions more consistent with power-law-like tails (Figure 2C–D; Supplementary Table 5). For cycles of length 5, *Streptococcus salivarius*, *Prevotella copri* Clade A, and *Faecalibacterium prausnitzii* continued to exhibit highly skewed overlap distributions consistent with log-normal behavior. In contrast, overlap distributions for *Oscillibacter* sp. ER4 and SGB15053 were comparatively less skewed, indicating that overlap between social and microbial cycles was more evenly distributed across individuals for these species (Figure 2C–D; Supplementary Table 5).

Together, these findings indicate that the degree of overlap between social and microbial cyclic structures varies substantially across microbial species and cycle lengths. For some species, overlap is concentrated among a relatively small subset of individuals, suggesting that social and microbial cyclic organization are closely aligned within particular social groups. For other species, overlap is more diffuse, indicating weaker correspondence between social and microbial cycle structure.

### Relationship between overlap and cycle participation

Building upon the observed overlap distributions, we next examined whether microbial species can be categorized into groups based on their overlap and cycle distributions. Conceptually, in a network, microbial species can vary along two dimensions: the extent of cycle participation and the degree of overlap with social cycles. Species with high cycle participation and high overlap may reflect transmission processes aligned with social interactions, whereas those with high cycle participation but low overlap may indicate transmission pathways less dependent on the social network. Our analysis revealed that most species form a generalized cluster characterized by low overlap and moderate cycle enrichment. Only a small subset of species exhibit strong deviations from this pattern, with some showing pronounced over-representation or under-representation of cycles, despite minimal overlap with social structure (Fig. S3). These outliers drive the observed heterogeneity and suggest species-specific transmission dynamics.

Next, we investigated whether cross-network overlap depends on an individual’s participation in social cycles, analyzing this relationship for each species separately. A negative-binomial regression model revealed a strong positive association among several species (Figure S4-5), indicating that increased participation in social network cycles is associated with higher overlap for these species: *Streptococcus salivarius* (SGB8007 group), *Prevotella copri Clade A* (SGB1626), *Faecalibacterium prausnitzii* (SGB15316 group), and SGB15053 (*Ruminococcaceae* family). This pattern suggests that, despite the overall negative association between microbial and social cycle participation, individuals participating in many social cycles were more likely to exhibit overlap with these microbial cycles.

## Discussion

Using strain-level microbial transmission networks and social networks from 1,787 individuals across 18 Honduran villages, we find that network cycles are a widespread motif in both microbial and social interaction systems. Across most microbial species, cycles were significantly over-represented relative to randomized networks preserving degree distributions, indicating that microbial transmission is not organized as a purely random or locally independent process. Instead, microbial circulation may occur within closed and recurrent groups of individuals. Similar motif enrichment was observed in the friendship networks, consistent with previous work demonstrating excess cyclic structure in social networks^15^. Together, these findings extend higher-order network analysis to microbial strain-sharing networks and suggest that repeated exposure within clustered groups may be an important feature of microbiome transmission dynamics.

At the individual level, participation in cycles was highly heterogeneous in both network types. Social cycle centrality exhibited strongly right-skewed distributions, with most individuals participating in relatively few cycles and a small subset participating in many. Similarly, many microbial species displayed heavy-tailed patterns, although the specific distributional forms varied across taxa. Some species, including *Faecalibacterium prausnitzii* and *Oscillibacter sp.* ER4, showed pronounced heavy-tailed cycle participation across different cycle lengths, whereas others were better approximated by log-normal behavior (like *Ruminococcaceae* SGB15053 sp.) or did not fit either model. These results indicate that microbial cyclic patterns are species-specific and that certain individuals may disproportionately contribute to the circulation of particular microbes.

The extent of cyclic organization also differed substantially across species. While most taxa showed a strong prevalence of cycles, a smaller subset (like *Catenibacterium sp. AM22 15*) exhibited depletion relative to null models. Importantly, these differences were not explained by prevalence alone. Species with comparable numbers of carriers often differed substantially in the density of strain-sharing ties, leading to variation in cycle counts. This suggests that cyclicity reflects not simply how common a species is within the population, but how transmission pathways are organized across hosts, and that these mechanisms likely depend on the biological traits of each microbe.

### Ecological Factors in Microbial Cyclicity

Specifically, several taxa with pronounced cyclic motifs were anaerobic organisms, including *Faecalibacterium prausnitzii* and *Roseburia inulinivorans*. Because many anaerobes survive poorly outside the host, their persistence is likely dependent on repeated short-range exposure through close contact or shared local environments. Cycles provide a natural structure for such repeated exposures by reinforcing circulation within connected individuals. In addition, some of these taxa participate in cross-feeding and other metabolic interactions that may further favor repeated transmission within such groups.^16,17^ In contrast, species with depleted cyclicity may rely more heavily on broader environmental dispersal or acquisition pathways that are less dependent on recurrent exposure.

### Independence and Divergence from Social Structure

Despite the widespread presence of cycles in both microbial and social networks, the relationship between the two was limited. Significant associations between microbial and social cycle participation were observed for only a small subset of these microbes, and several of these associations were negative. In particular, *Streptococcus salivarius* and SGB1883 of the *Bacteroidaceae* family showed negative associations between microbial and social cycle centrality across all cycle lengths. In addition, species like *Oscillibacter sp. ER4*, unknown species of the *Ruminococcaceae* family (SGB15053 and SGB15085), and *Candidatus Gastranaerophilales bacterium* showed negative associations for some cycle lengths. These findings suggest that individuals embedded in highly cyclic microbial transmission structures are not necessarily those occupying highly cyclic positions in the social network.

For several of these taxa, ecological characteristics may help explain this pattern. Streptococcus salivarius is primarily an oral organism and may be transmitted through environmental exposure, diet, or aerosolized droplets, meaning it does not rely solely on close social contact^19,20,24^.

Similarly, *Oscillibacter sp. ER4* and several uncharacterized *Ruminococcaceae* taxa are anaerobes strongly associated with host dietary and physiological factors^22,23^. Their weaker alignment with social cycle structure suggests that microbial persistence may depend on localized ecological conditions, shared environments, or host-specific factors that are only partially captured by friendship networks. More broadly, these results support the idea that microbial transmission networks can form niche-like structures that operate – at least, partly – independent of the social interactions.

### Deconstructing Cycle Overlap versus Centrality

The overlap analyses further highlight this partial independence between social and microbial cyclic organization. Although some individuals participated in overlapping social and microbial cycles, the extent of overlap varied across species and cycle lengths. For certain taxa, overlap was concentrated among a small number of individuals, whereas for others it was distributed more evenly throughout the population. At the species level, most microbes occupied regions of relatively low overlap despite substantial cycle enrichment, indicating that strongly cyclic microbial transmission does not necessarily require alignment with social cycles. On the other hand, a smaller number of outlier species exhibited comparatively high overlap (like the unknown sp. SGB13947 from the *Firmicutes* phylum), suggesting closer alignment between microbial circulation and social networks.

Importantly, the overlap analyses capture a distinct property of the system from the association models. Cycle centrality measures how strongly individuals participate in cyclic structures within a single network, whereas overlap measures whether the same individuals participate in cycles across both network types. Consistent with this distinction, we found that individuals participating in many social cycles were often more likely to exhibit overlap between social and microbial cycles, even when microbial cycle centrality itself was negatively associated with social cycle participation. Together, these findings suggest that microbial transmission can simultaneously exhibit partial dependence on social structure while also maintaining species-specific pathways that are relatively independent of it.

### Limitations and Future Directions

This study has several limitations. First, strain-sharing cannot completely distinguish direct interpersonal transmission from acquisition through shared environments or unobserved contacts. Second, strain inference relies on currently available metagenomic methods and does not provide complete strain characterization. Third, because transmission directionality is unavailable, all analyses were conducted using undirected networks. Future work incorporating temporal sampling, environmental measurements, and directional transmission models may help clarify the mechanisms underlying cyclic microbial circulation.

Overall, our findings demonstrate that microbial strain-sharing networks exhibit rich higher-order organization that only partially mirrors human social structure. Network cycles provide a useful framework for studying repeated exposure, localized persistence, and heterogeneous transmission pathways across microbial species. These results suggest that understanding microbiome transmission may require moving beyond pairwise interactions toward higher-order motifs that capture specific recurrent patterns of host contact and exposure.

## Acknowledgments

We thank all the study participants in Honduras. We thank Jose Eduardo Gámez for coordinating the field work; and Rennie Negron, Liza Nicoll, and Thomas Keegan for their support on field operations, data collection, and administrative support.

This work was supported by the NOMIS Foundation, with additional support from the Pershing Square Foundation and Paul Graham. Empanelment of the underlying cohort was supported by the Bill and Melinda Gates Foundation.

## Author contributions

Conceptualization: SVS, GI, and NAC; Methodology: SVS and NAC; Analyses: SVS; Funding acquisition: NAC; Writing: SVS, FB, GI, YC, and NAC.

## Data and Code availability

The full datasets generated and/or analyzed during the current study is not publicly available due to privacy concerns. However, much of the data regarding the microbes and social ties is already available at ZENODO links. The code for replicating this analysis is available at https://github.com/human-nature-lab/socio-microbial_strain_cycles.

Metagenomic data is available: https://www.ncbi.nlm.nih.gov/bioproject/PRJNA999635.

Abundance tables and certain strain-level information are available here: https://zenodo.org/records/11150476. Core metadata for each subject (their age, sex, BMI, Bristol Stool Scale, and village ID) is publicly available here: https://zenodo.org/records/11150476. Additional, more confidential metadata (as specified by human subjects constraints) are available in two separate files, and are archived at https://zenodo.org/records/11153185 and at https://zenodo.org/records/11153210.

## Methods

Participants resided in 18 isolated villages in the western highlands of Honduras. The average distance from each of the 18 villages to the nearest other village among the is 1.1km, and the average distance to the farthest other village is 24.7km. These 18 villages range in size from 66 to 432 individuals, and their underlying average household size is 3.49. The average age of participants is 41 (SD=17; range: 15-93); 62% are women; and 41.8% are married.

### Social network construction

Village based social networks were constructed based on responses to a hour-long survey of 1,787 adults and adolescents (age > 15) across the 18 isolated villages, using photographic survey tools^25^. The survey was implemented in Trellis^26^, and individuals were asked to identify the family members, friends, and other sorts of perceived relationships using name generators. For the purpose of this work, we generated an undirected social network using all friendship connections between individuals living in the same village and in different households.

The Yale IRB and the Honduran Ministry of Health approved all data collection procedures (Protocol #1506016012). All participants provided informed consent before enrolling in the study. Also, all our methods are approved and performed under guidelines by Yale Committee on Human Subjects.

### Sample preparation

All participants were asked to self-collect fecal sample by following instructions on a training module, which was later stored in a -80 C freezer, before being transported to the United States, where it was sequenced. Further details on the sample preparation and sequencing procedure can be found in previous work^7^.

### Strain characterization and microbial network construction

Metagenomes were preprocessed using prinseq, BMTagger, and Trimmomatic^27^. Species-level profiles were generated by MetaPhlAn 4^3^. We identified a total of 2,508 species, of which 639 species were retained after filtering for low abundance at the individual level or low prevalence at the population level. Strain-level phylogenies were generated by StrainPhlAn^3^;using species-based threshold and the strain_tramission.py script provided by StrainPhlAn, we identified pairs of individuals sharing the same strain. Strain-sharing connections were then used to create an undirected microbial network. That is, ties between individuals living in the same village but in different households were used for constructing a microbial sharing network.

### Power-law distribution

All our power-law model fits and evaluation were performed using the R package *poweRlaw*^28^(with function “conpl”).

### Cycle construction in social and microbial networks

Using prior modelling approaches as a foundation^18,29^, we defined cycles *C_uv_* as a path where the first and last node coincide (are identical). We define the set of all cycles of length *k*, of which node *n* is a member, as follows:

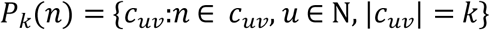

and the set of all cycles of length *k* in the network graph as:

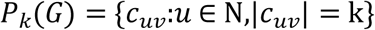

We use the *k*-cycle metric from previous work^15^, which is defined as the number of network cycles of length *k* that a node belongs to:

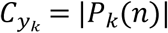

We present the exact formulation used in our previous work^15^ in the cycle centrality metric, or the number of cycles each individual participates in at the individual level, for cycles of length ∈ {3,4,5}. These are estimated using SNA^30^ package in R, across friendship networks (social networks) for each village, and strain-sharing networks (microbial networks) per species and village. Therefore, in total we have 18 villages across 639 species totaling 11,502 distinct microbial networks for analysis. For our regression models, we use 639 species networks for exploring and characterizing relationships.

After computing cycles, the number of cycles each person participates in was generated using above equations, for each individual across both social and microbial networks. The full list of cycles prevalence per species can be found in Supplementary Table 1.

### Overlap of cycles across social and strain-sharing networks

We designed a metric analogous to the Jaccard index to detect cycle overlap across modalities in the same set of nodes (*N*). We define two node sets *M* and *S*, such that *M, S* ⊆ N, where *M* and *S* are nodes belonging to microbial and social network in the same network comprising of *N* nodes (superset).

At the same time, the cycles a person *i* participates in can be defined as:

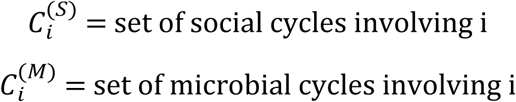

Among the cycles, the nodes involved in cycles across modalities can be defined by:

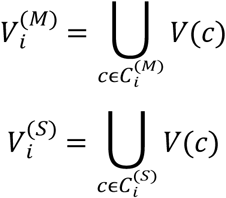

Where V(c) are the individuals participating in cycle c, where V(c)∉i.

The multi-modal *overlap ratio* can be defined as the individuals common to both sets, or,

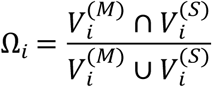

This overlap ratio can range from 0 (no overlap) to 1(full overlap).

### Regression-based statistical model

We implemented a mixed-effects beta regression framework to quantify the association between social cycle centrality and microbial cycle centrality for each species across 1,787 individuals.

The response variable was defined as the per-individual microbial cyclic centrality, computed as the fraction of cycles in which an individual participates within a given village and species-specific network.

Because beta regression requires response variables to lie strictly within the open interval (0,1), values of 0 and 1 were adjusted using a standard transformation (e.g., Smithson and Verkuilen, 2006^31^). We used a logit link function to model the mean of the beta distribution.

The model specification was:

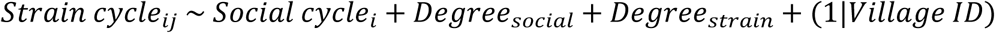

where social and microbial degree terms were included to control for individual-level connectivity in each network. A random intercept for village was included to account for clustering and variation in network structure across villages.

Regression coefficients were computed from the association models specific to each species and cycle length. To account for multiple hypothesis testing across species, we applied the Benjamini–Hochberg procedure to control the false discovery rate, yielding adjusted p-values for each species, yielding an adjusted P-value for every species in the mixed regression model. ^32^

Beta regressions were perfomed using “glmmTMB” function from the “glmmTMB” package (v1.1.4)^33^ with the family parameter of “beta_family”.

A negative binomial regression-based framework was used to compute the association between overlap of microbial and social cycles with participation in social cycles, using the following equation:

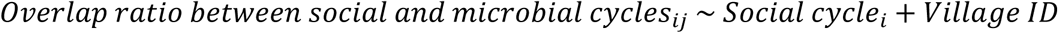

These overlap values for every individual, similar to social cycle centrality per individual, was used to measure the association separately for every species and cycle length {3,4,5}. As before, all P-values were corrected for Benjamini Hochberg procedure. All associations from the regression model are available in Supplementary Table 4.

Negative binomial regression models were fit using the glmer.nb function from the lme4 package (v1.1-35.3) to model count-based outcomes, while beta regression models were used for normalized cycle participation measures, as described above.

Linear paths were not included as covariates due to strong collinearity with cycle-based measures in dense networks.

### Null permutation models

Null permutation models were used for validating cyclicity in microbial networks and the association between social and microbial cycle participation. Null permutation models are primarily used to compare how different the real value is compared to values originating from randomized villages networks. For this purpose, we created null distribution of villages (N=1,000) with the same network structures (degree and edge distribution) and the number of individuals within a village, similar to our previous work^15^. After creating null villages, we computed the number of cycles, and the relationship between social cycle centrality and microbial cycle centrality per person. Cycles generated in both real and null-permuted networks (*N* = 1,000) were compared using Z-scores:

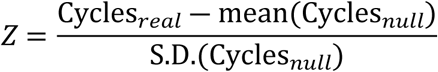

Strictly, Z > 0 indicated over-representation of cycles and Z < 0 indicated under-representation of cycles in real networks compared to null networks.

All associations from the null permutation model are available in Supplementary Table 3.

---

**Figure S1:**
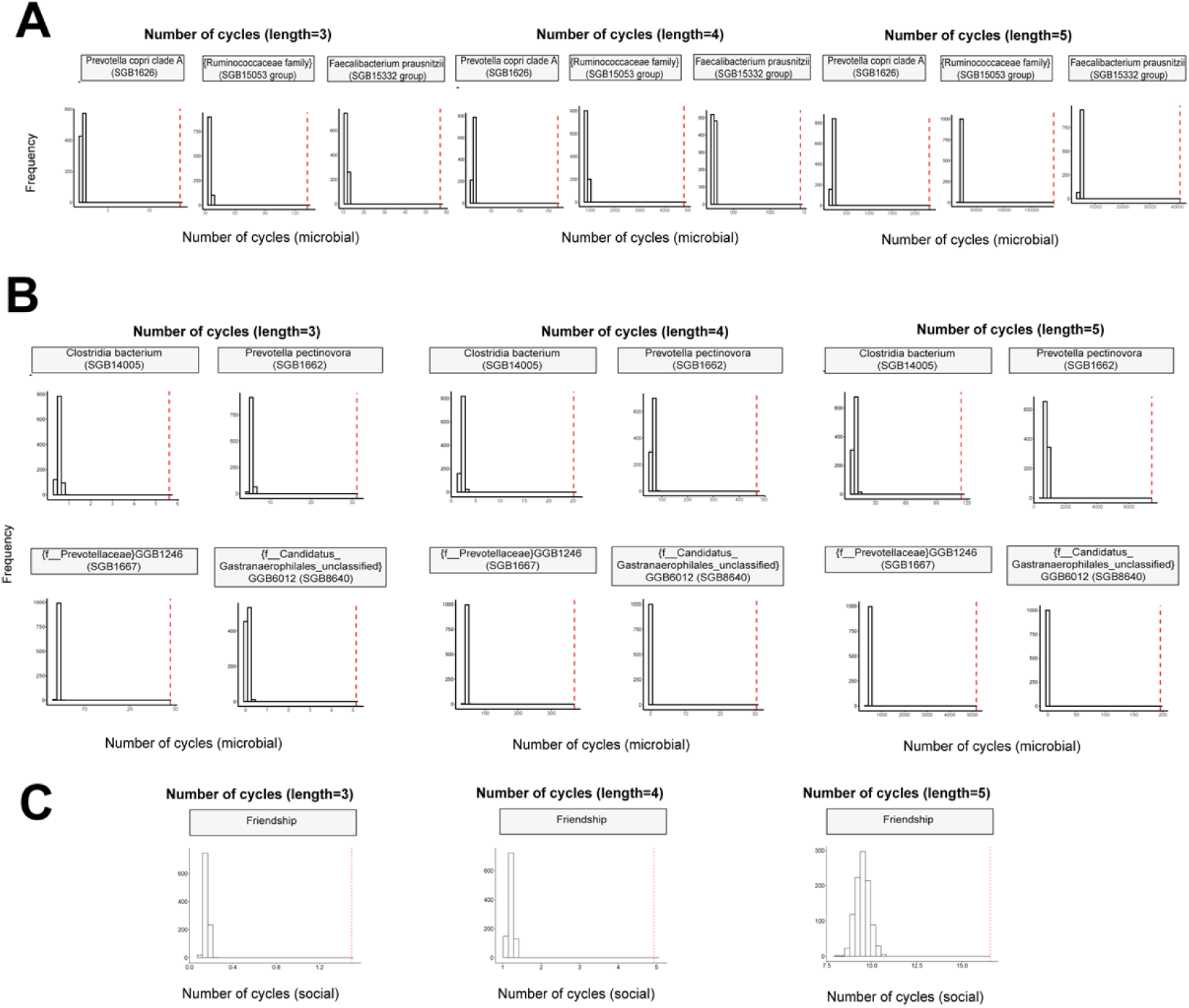
Histogram of average number of cycles detected. Red vertical dotted line indicates average detected per person from real villages. This process was repeated for null permuted villages (N=1000) and for all cycle lengths (lengths ∈ {3,4,5}). This plot demonstrates that observed cycles is different from cycles emerging from expected or random villages (p-value<10^-25^).

**Figure S2:**
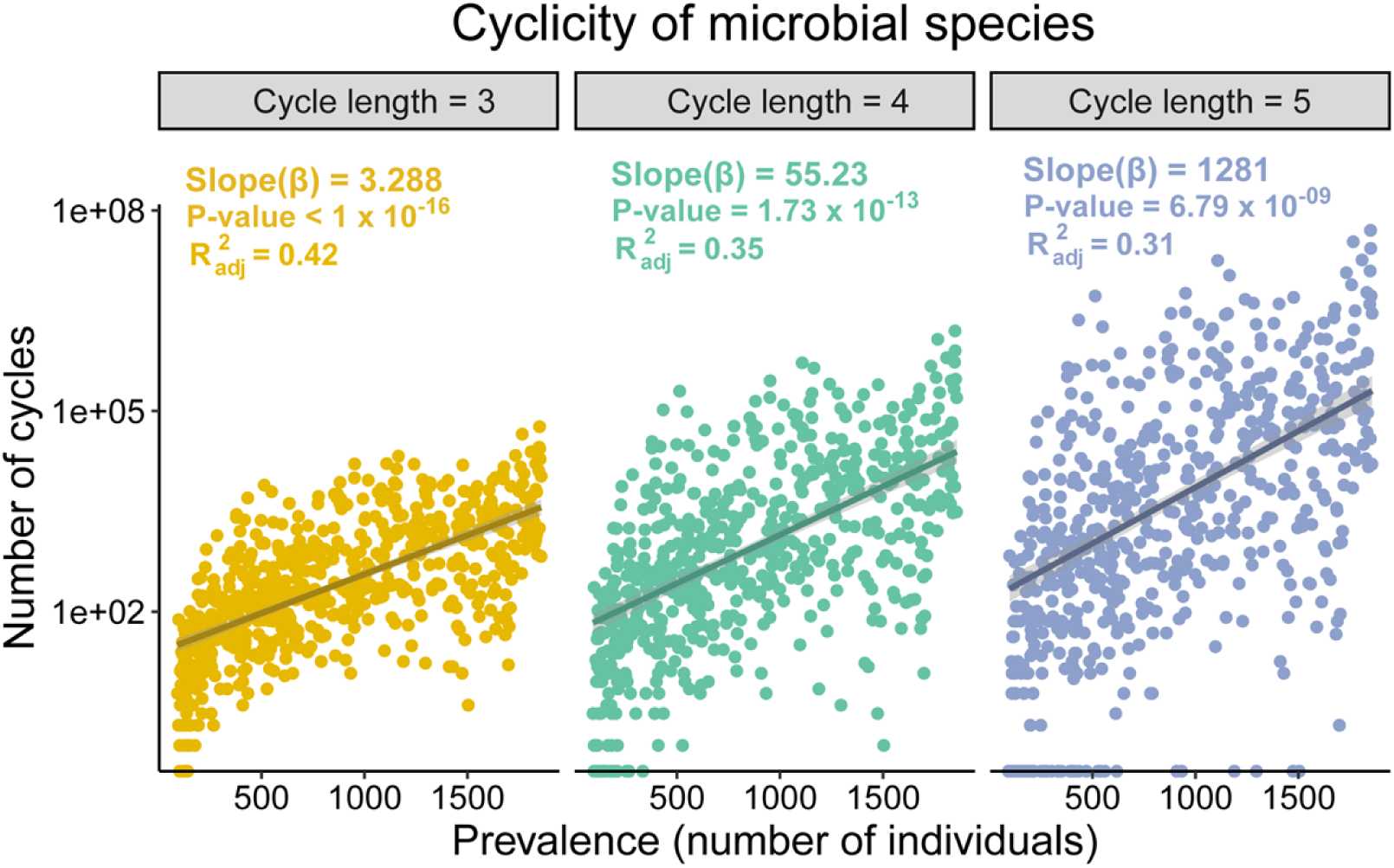
Cyclicity of species. Total number of cycles for every species against the number of people each species is prevalent in. The trend lines shows how the number of cycles varies with prevalence in a linear model and the corresponding slopes, p-values, and R^2^ (goodness of fit) of the trend. Every dot is a unique species. This plot contains all 639 species.

**Figure S3:**
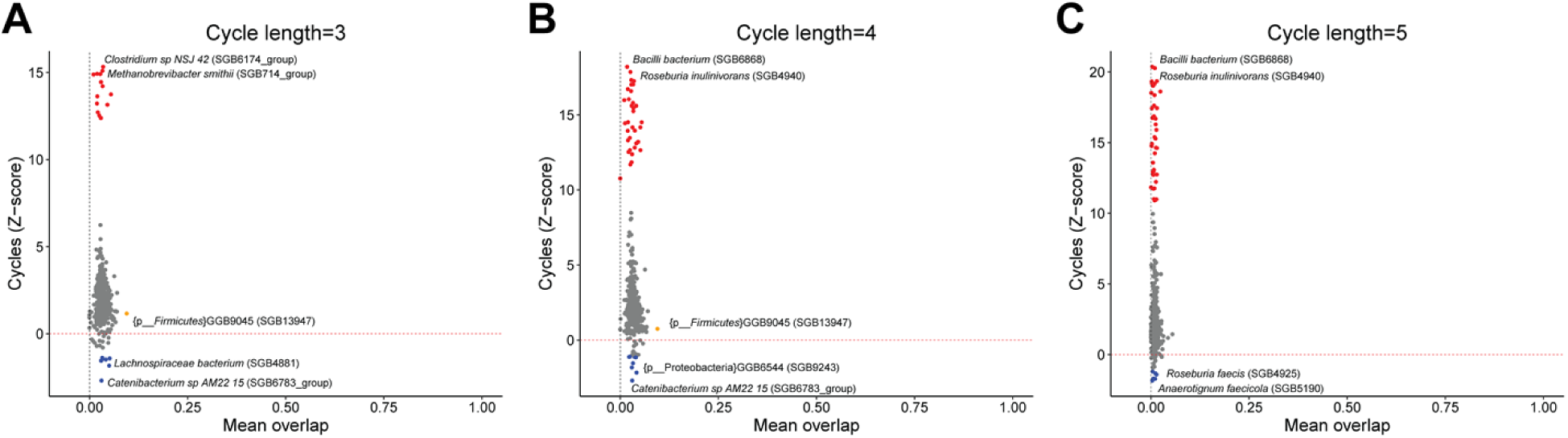
Relationship between Cyclicity and Overlap. The relative comparison of microbial cycles (against randomized networks) of all lengths and species were plotted against their overlap with social cycles. All three plots show a consistent trend – most species aggregate in the low-overlap and low-overrepresented region, with few species separating from this trend. The top two outlier species in each direction are labeled. Red, blue, and orange dots depict species with highly over-represented cycles, under-represented cycles, and overlap respectively.

**Figure S4:**
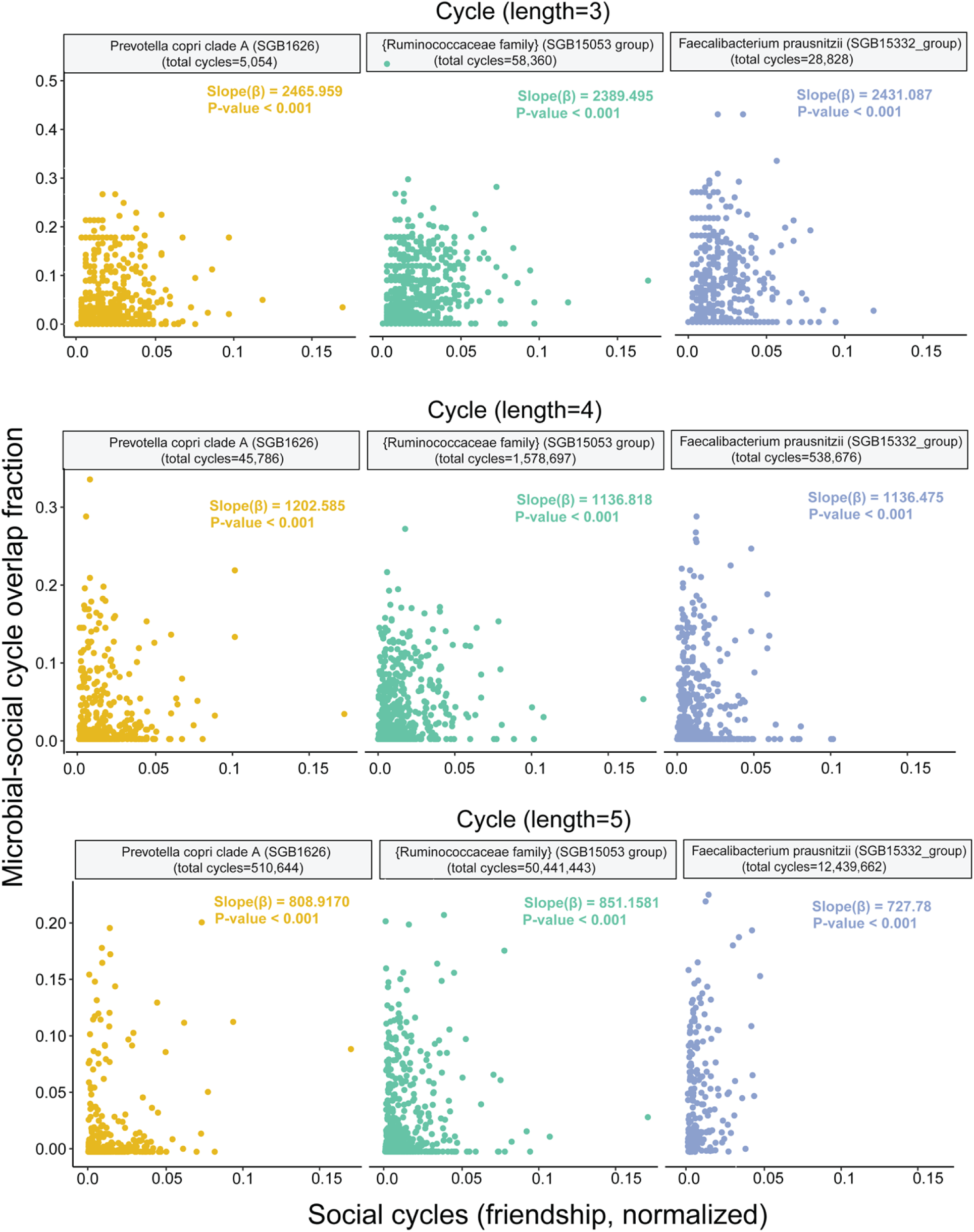
Association of overlap with social cycle. Negative-binomial regression plots for association between overlap fractions (between microbial and social cycles) and social cycle participation for cycles of lengths {3,4,5}.

**Figure S5:**
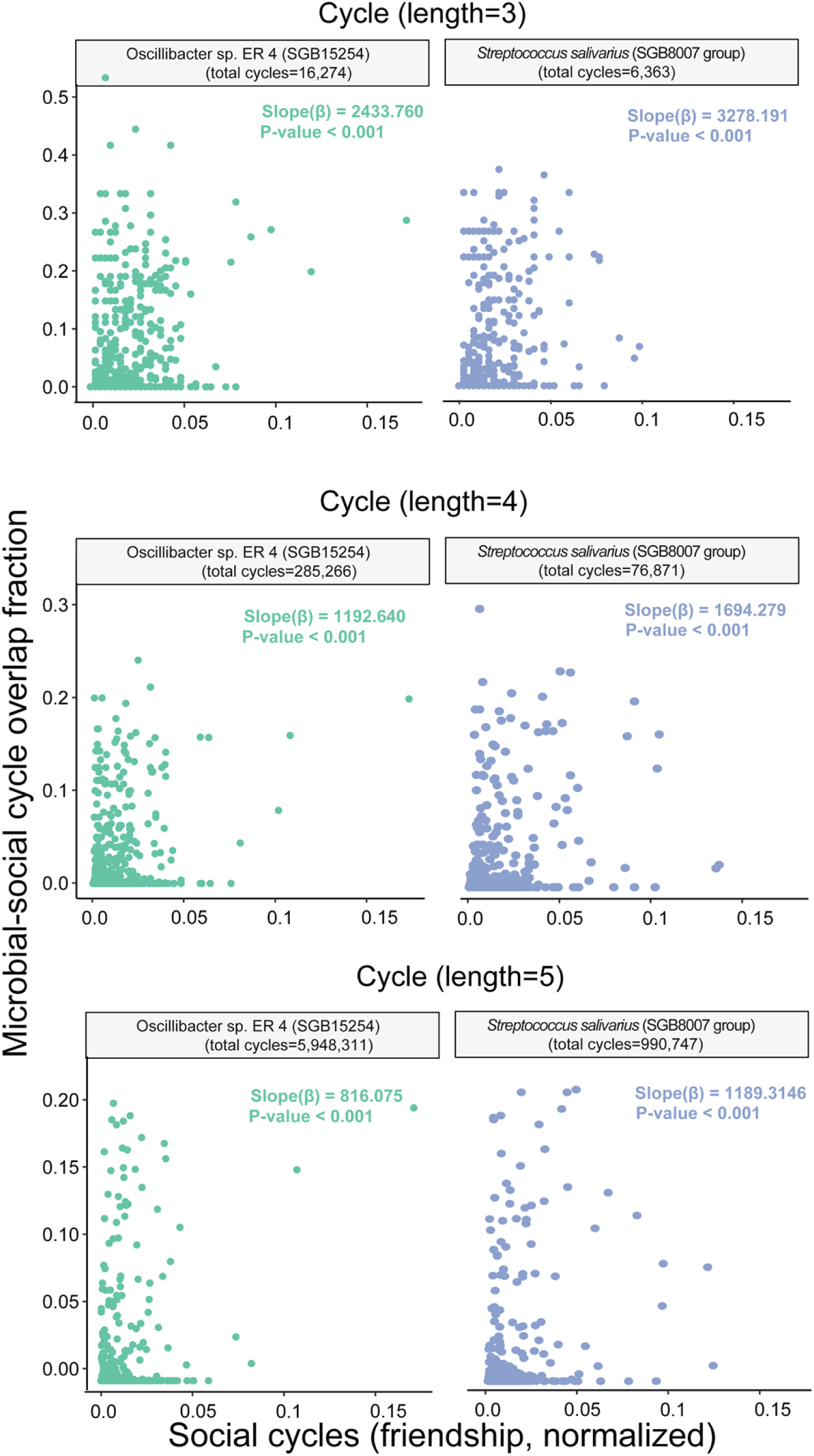
Association of overlap with social cycle for significant species (from association) Negative-binomial regression plots for association between overlap fractions (between microbial and social cycles) and social cycle participation for cycles of all lengths {3,4,5}.

